# Electrophysiological Comparison of Mechanosensitive PIEZO1 Channel Currents across Different Human Cell Types

**DOI:** 10.64898/2026.11.20.768008

**Authors:** Katsuhiko Muraki

## Abstract

The detection of mechanical stress is essential for a wide range of physiological functions in humans, including touch sensation, hearing, and the maintenance of vascular myogenic tone. PIEZO1 is a mechanosensitive cation channel that is widely expressed in both neuronal and non-neuronal cells and plays critical physiological roles in multiple organs. However, the electrophysiological comparisons of endogenous PIEZO1-mediated mechanosensitive currents have remained limited across different human cell types under identical experimental conditions. Here, the author compared the electrophysiological properties of endogenous human PIEZO1 (hPIEZO1) currents in four cultured human cell types: human aortic endothelial cells, human brain capillary endothelial cells, human fibroblast-like synoviocytes, and human osteoblast-like MG63 cells. Under whole-cell patch-clamp conditions, application of the PIEZO1 agonist Yoda1 robustly activated hPIEZO1-like currents in all four cell types. In excised outside-out patch recordings, application of positive pressure to membrane patches from these cells also elicited PIEZO1-like channel currents, which were markedly potentiated by Yoda1 and effectively inhibited by gadolinium, a non-selective cation channel blocker. Furthermore, the unitary current amplitudes of single channel events evoked by mechanical stimulation were comparable to those elicited by Yoda1 across all four cell types. Collectively, these findings provide electrophysiological evidence that endogenous PIEZO1 functions as a mechanosensor in diverse human cell types and suggest that PIEZO1 contributes to mechanically regulated physiological processes in multiple human organs.

## 1. Introduction

Mechanical stress is a fundamental physiological stimulus that underlies a wide range of biological functions, including touch sensation, hearing, and the maintenance of vascular myogenic tone. In the vascular system, shear stress exerting on endothelial cells activates multiple ion channels, thereby modulating vascular function. Accordingly, endothelial mechanosensors play critical roles in the regulations of blood pressure, arteriosclerosis, and vascular remodeling. Consistent with this concept, the mechanosensitive cation channel PIEZO1, originally identified as a mechanically activated ion channel [1, 2], is highly expressed in vascular endothelial cells across multiple species and plays essential roles in vascular development and the regulation of vascular tone [3, 4]. Since the discovery of PIEZO family, extensive studies have demonstrated that PIEZO channels, PIEZO1 and PIEZO2, are evolutionally conserved across vertebrates and invertebrates and are also present in plants and protozoa, suggesting that PIEZO-mediated mechanosensation represents fundamental biological functions [1].

PIEZO channels are directly activated by a variety of mechanical stimuli, including membrane stretch, shear stress generated by physiological fluid flow, membrane indentation by a probe, and changes in membrane stiffness [1, 3–5]. Their roles in mechanotransduction have been extensively studied using a variety of animal models, including as mouse [3, 4], zebrafish [6], and *drosophila* [7, 8]. Notably, the discovery of Yoda1 (2-[5-[[(2,6-Dichlorophenyl)methyl]thio]-1,3,4-thiadiazol-2-yl]-pyrazine), a synthetic PIEZO1 agonist [9], has greatly facilitated investigation of the diverse physiological functions of PIEZO1. These functions include sensing bladder distention in the urothelium [10], mechanical stress-induced osteoclastogenesis [11], and regulation of red blood cell function [12–15].

Despite substantial advances in the biophysical, structural, physiological, and pathphysiological characterization of PIEZO1, direct comparisons of the electrophysiological properties of endogenous PIEZO1 across different human cell types remain limited. Studies of other ion channels have demonstrated that the functional properties and physiological roles of a given ion channel can vary substantially among tissues and organs. In the vascular endothelium, shear stress-induced activation of transient receptor potential vanilloid 4 (TRPV4) channels promotes nitric oxide production, thereby contributing to vascular smooth muscle relaxation [16]. In contrast, TRPV4 contributes to serotonin-induced pulmonary vasoconstriction in chronic hypoxic pulmonary hypertension [17]. Such tissue- and context-dependent functions of ion channels highlight the importance of characterizing their properties in different cellular environments and may provide a basis for the development of tissue- or organ-targeted pharmacological interventions. Therefore, determining whether the electrophysiological and pharmacological properties of human PIEZO1 (hPIEZO1) differ among human cell types is important for understanding its tissue-specific physiological functions and for evaluating its potential as a therapeutic target.

Endogenous PIEZO1 has been shown to be functionally expressed in a variety of humans cell types and tissues, including vascular endothelial cells [3, 4], red blood cells [18, 19], osteoblasts [20], fibroblasts [21], and neurons [1, 22]. However, the electrophysiological properties of these endogenous PIEZO1s have not been compared under identical experimental conditions. In the present study, the author therefore compared the electrophysiological characteristics of endogenous hPIEZO1 in four cultured human cell types: commercially available human aortic endothelial cells (AORET), human brain capillary endothelial cells (BCET), human fibroblast-like synoviocytes (FLS), and human osteoblast-like MG63 cells (MG63). Using whole-cell and excised outside-out patch-clamp recordings, we examined Yoda1-induced and mechanically activated hPIEZO1 currents under comparable experimental conditions. The present results demonstrate that the fundamental electrophysiological properties of endogenous hPIEZO1 are largely conserved across these four human cell types, including their responses to mechanical stimulation and Yoda1.

## 2. Results

### 2.1. Effects of Yoda1 on endogenous hPIEZO1 currents under whole-cell recording conditions

The author first examined the effects of Yoda1 on endogenous hPIEZO1 currents in human aortic endothelial cells (AORET), human brain capillary endothelial cells (BCET), human fibroblast-like synoviocytes (FLS), and human osteoblast-like MG63 cells (MG63). As shown in Figures 1A-1D, under whole-cell recording conditions, application of 3 μM Yoda1 reversibly activated PIEZO1-like currents in all four human cell types. Quantitative analysis confirmed that Yoda1 significantly increased the current amplitude in each cell type (Figure 1E), indicating the functional expression of Yoda1-sensitive PIEZO1 channels. The amplitude of Yoda1-induced currents measured at -90 mV did not differ significantly among the four cell types (Supplementary Figure S1A).

**Figure 1.**
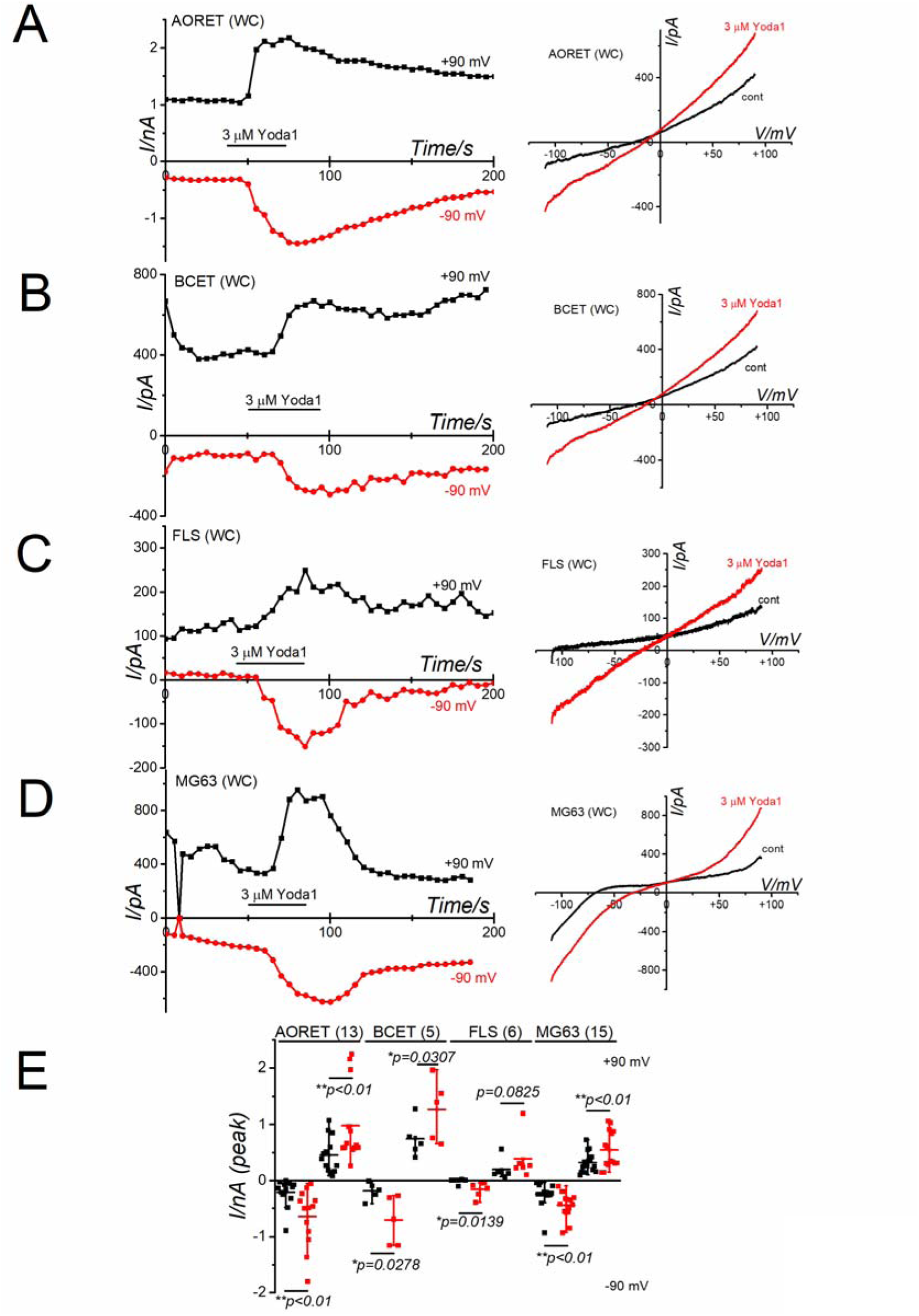
Effects of Yoda1 on endogenous hPIEZO1 currents under whole-cell recording conditions. (A-D) Human aortic endothelial cells (AORET; A), human brain capillary endothelial cells (BCET; B), human fibroblast-like synoviocytes (FLS; C), and human osteoblast-like MG63 cells (MG63; D) were voltage-clamped in the whole-cell configuration and exposed to 3 μM Yoda1. Left panels: Voltage-ramp pulses from −110 to +90 mV for 300 ms were applied every 5 s, and the current amplitudes measured at −90 and +90 mV were plotted against time. Arrows denote the time points at which the corresponding current-voltage (I-V) relationships were obtained. Right panels: Representative I-V relationships obtained before and during application of 3 μM Yoda1. (E) Summary of current amplitudes at -90 and +90 mV in the absence and presence of 3 μM Yoda1 in AORET, BCET, FLS, and MG63 cells (n=13, 5, 6, and 15 independent experiments, respectively). Statistical significance was assessed using paired Student’s *t*-tests.

### 2.2. Mechanical activation of endogenous hPIEZO1 currents in excised outside-out patches

To characterize the mechanical sensitivity of endogenous hPIEZO1, positive pressures ranging from 2.5 to 100 mmHg was applied to excised outside-out membrane patches from each human cell type. Although the magnitude of pressure-induced current activation varied from cell to cell, robust responses were observed in AORET and BCET cells (Figures 2A and 2B). Pressure-induced currents were also observed in FLS cells and MG63 cells (Figures 2C and 2D), whereas the responses in all four human cells were similar (Supplementary Figure S1B). To compare the mechanical sensitivity of endogenous hPIEZO1 among the four cell types, pressure-response relationships obtained over the range 2.5-100 mmHg were fitted with the Boltzmann equation (right panels of Figures 2A-2D). Satisfactory Boltzmann fits were obtained for 12 out of 19 patches form AORET cells, 4 out of 10 patches from BCET cells, 3 out of 8 patches from FLS cells, and 4 out of 23 patches from MG63 cells. Among the successfully fitted datasets, neither midpoint pressure (p50) nor the slope coefficient (dp) differed significantly among the four cell types (Supplementary Table S1). These results indicate that mechanically activated hPIEZO1-like currents are present in all four human cell types and that their estimated pressure sensitivity is comparable under the present experimental conditions.

**Figure 2.**
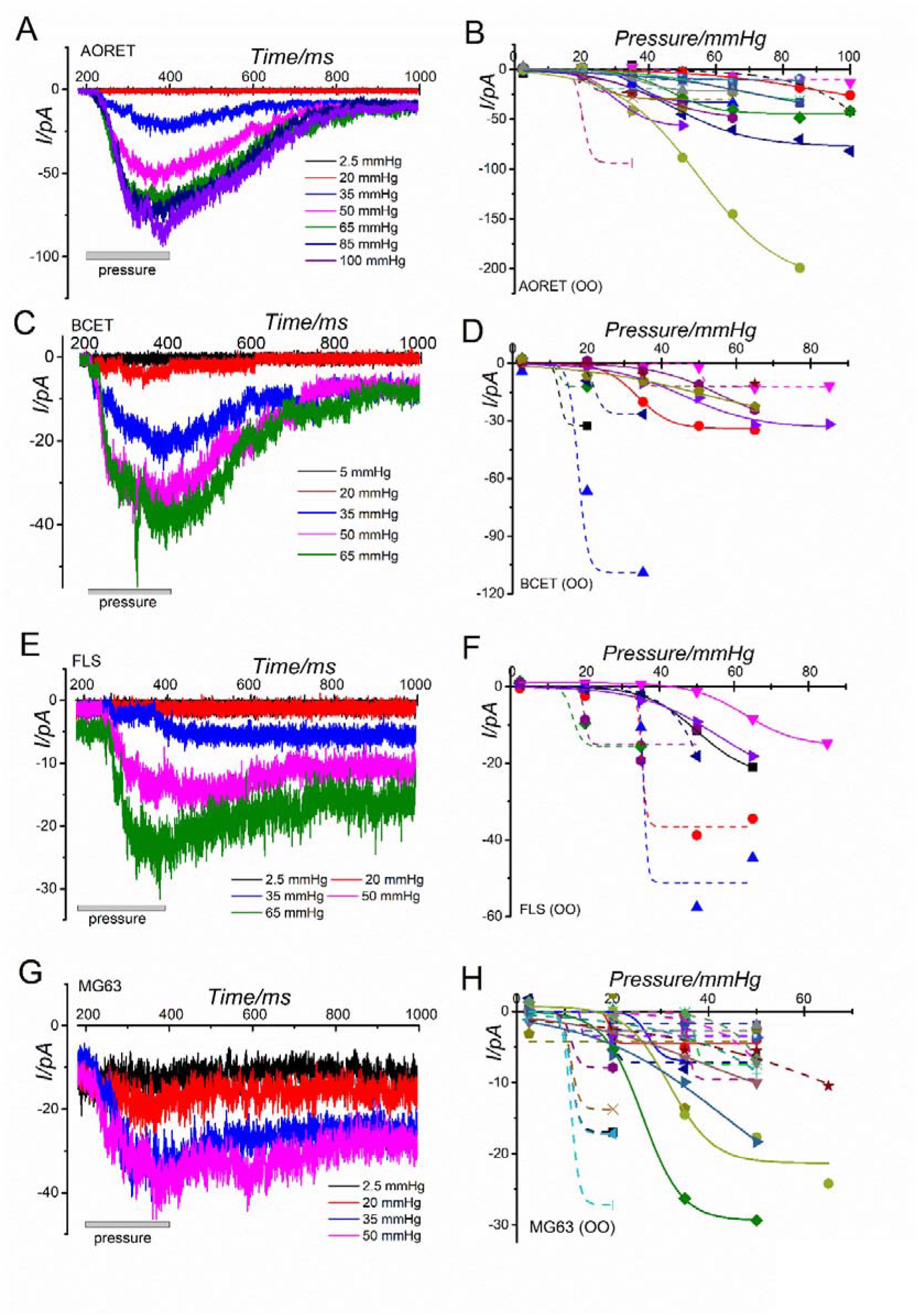
Pressure-induced membrane currents in human cells under excised outside-out patch conditions. (A-H) Excised outside-out membrane patches from AORET (A), BCET (C), FLS (E), and MG63 (G) were voltage-clamped and subjected to positive pressures from 2.5 to 100 mmHg. Left panels: Voltage-ramp pulses from −110 to +90 mV for 150 ms were applied at a holding potential of −60 mV, followed by application of positive pressures for 200 ms every 10 s. Right panels: Pressure-response relationships were examined using datasets obtained over pressure ranges of 2.5-100 mmHg for AORET (n=19, B), 2.5-85 mmHg for BCET (n=10, D), 2.5-85 mmHg for FLS (n=8, F), and 2.5-50 mmHg for MG63 (n=23, H). Peak pressure-induced currents were plotted against applied pressure and fitted with the Boltzmann function. Datasets for which satisfactory Boltzmann fits were obtained are shown as solid lines (12 out of 19, 4 out of 10, 3 out of 8, and 4 out of 23 patches for AORET, BCET, FLS, and MG63, respectively). The Boltzmann parameters, midpoint pressure (p50) and slope coefficient (dp), were summarized in Supplementary Table S1. Datasets that could not be satisfactorily fitted, including those containing only two data points, are shown as dushed lines.

### 2.3. Effects of Yoda1 and Gd^3+^ on mechanically activated endogenous hPIEZO1 currents

TRPV4, another channel implicated in mechanosensation, is functionally expressed in human endothelial cells [23]. We have also previously demonstrated functional expression of TRPV4 in human FLS cells and its potential involvement in immune regulation in human knee-joints [24]. Therefore, to further determine whether the pressure-induced currents observed in the present study were mediated predominantly by endogenous hPIEZO1, the author examined the effects of Yoda1 and Gd^3+^ on mechanically activated currents (Figures 3-5). Channel activity evoked by a positive pressure of 20 mmHg was significantly potentiated by 3 μM Yoda1 in all four human cell types (left panels of Figures 3A-3D). Moreover, discrete single-channel events were observed at -50 mV in some excised outside-out patches during mechanical stimulation and in the presence of Yoda1 (middle panels of Figures 3A-3D). The amplitudes of unitary currents induced by mechanical stimulation were compared with those induced by Yoda1 using current-amplitude histograms (right panels of Figures 3A-3D). The unitary current amplitudes evoked by the two stimuli were comparable (Figure 4 and Supplementary Table S2), consistent with the interpretation that mechanical stimulation and Yoda1 activate the same or similar populations of endogenous hPIEZO1 channels. The author next examined the effects of 3-10 μM Gd^3+^ on currents induced by 20 mmHg positive pressure (Figures 5A-D). The effects of Gd^3+^ and 3 μM Yoda1 on pressure-induced currents are summarized in Figures 5E-H. In all four human cell types, pressure-induced currents were significantly potentiated by Yoda1 and markedly inhibited by Gd^3+^. Together, these pharmacological findings support the conclusion that Yoda1-sensitive endogenous hPIEZO1 channels constitute a major component of the mechanically activated currents observed under these experimental conditions.

**Figure 3.**
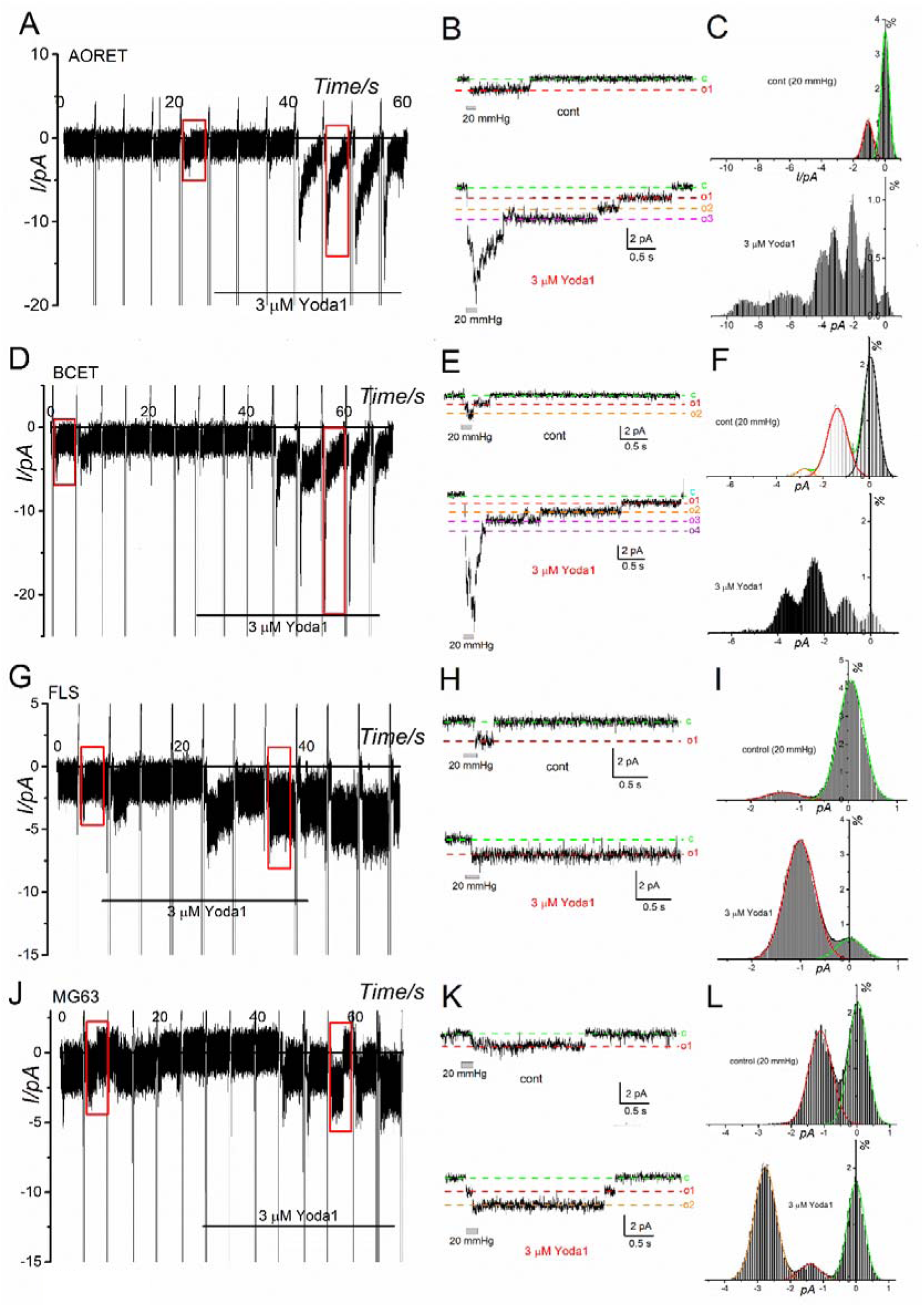
Effects of Yoda1 on pressure-induced currents in human cells under the excised outside-out patch conditions. Excised outside-out patches were subjected to a positive pressure of 20 mmHg in the absence and presence of 3 μM Yoda1. Voltage-ramp pulses from −100 to +100 mV for 150 ms were applied from a holding potential of −50 mV every 5 s. Left panels: Representative time courses of holding currents at −50 mV in AORET (A), BCET (D), FLS (G), and MG63 (J) cells. Each vertical deflection indicates application of a voltage-ramp pulse. Middle panels: Expanded current traces recorded in the absence and presence of 20 mmHg pressure before (upper traces in B, E, H, and K) and during (lower traces in B, E, H, and K) application of 3 μM Yoda1. Current signals were digitally filtered at 1 kHz. C and O1, O2, O3, and O4 denote the closed state and open states corresponding to one-, two-, three-, and four open channels, respectively. Right panels: Current-amplitude histograms constructed from 4-s recording period following pressure stimulation, as indicated by the red boxes in the left panels, in the absence (upper panels in C, F, I, and L) and presence (lower panel in C, F, I, and L) of Yoda1. Gaussian distributions were fitted to the amplitude histograms and superimposed on the data. The peak amplitude of the Gaussian components is summarized in Figure 4.

**Figure 4.**
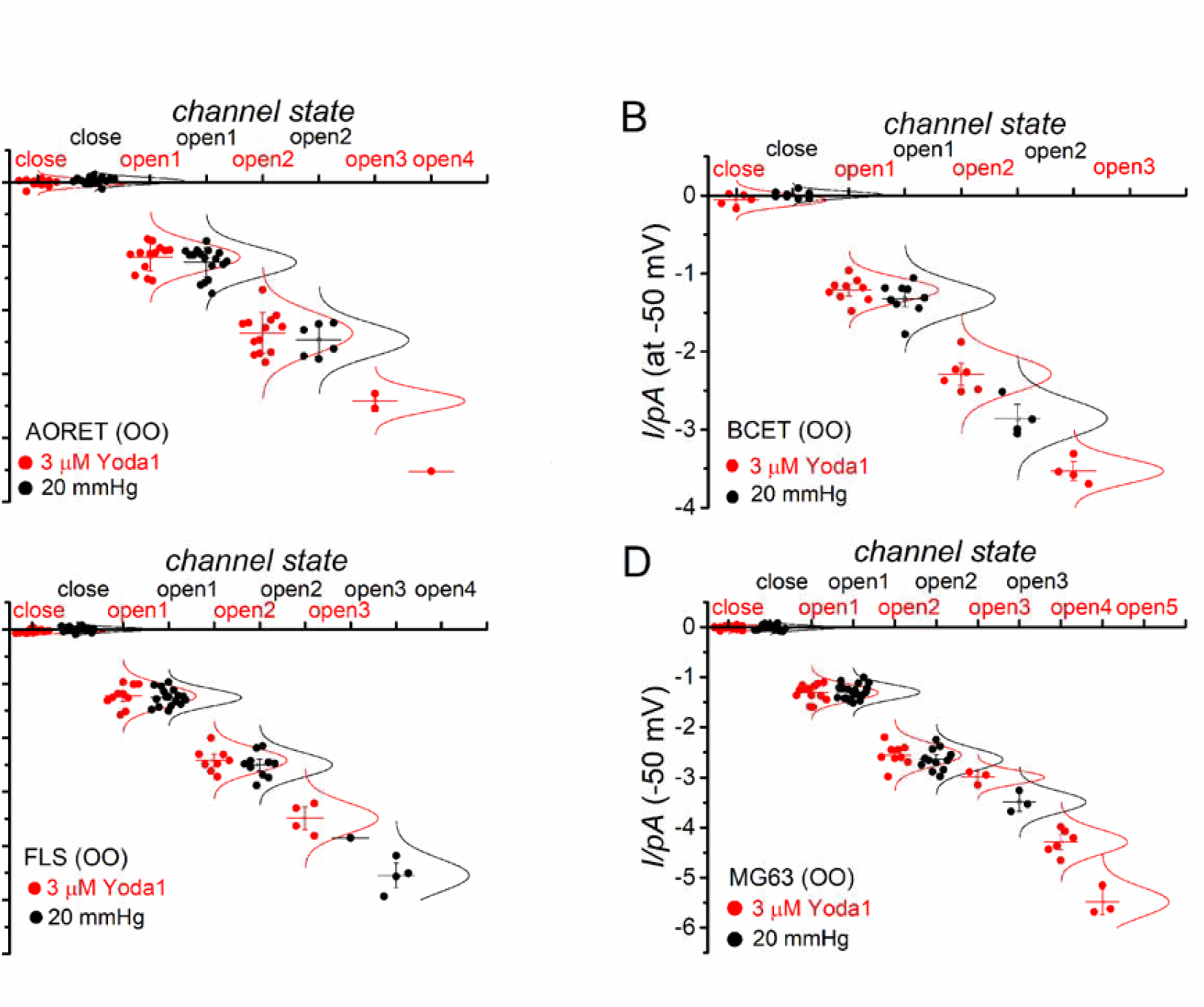
Comparison of pressure- and Yoda1-induced unitary currents in human cells. (A-D) Unitary current amplitudes at -50 mV were compared between channels activated by 20 mmHg positive pressure and those activated by 3 μM Yoda1. The labels “Close”, “Open1”, “Open2”, “Open3”, “Open4”, and “Open5” indicate the closed state and successive current levels corresponding to one through five open channels, respectively. The Open3 level observed in the presence of Yoda1 in MG63 cells may include a subconductance level of the Open2 state (D, Supplementary Table S2). Parameters obtained from Gaussian fitting are summarized in Supplementary Table S2.

**Figure 5.**
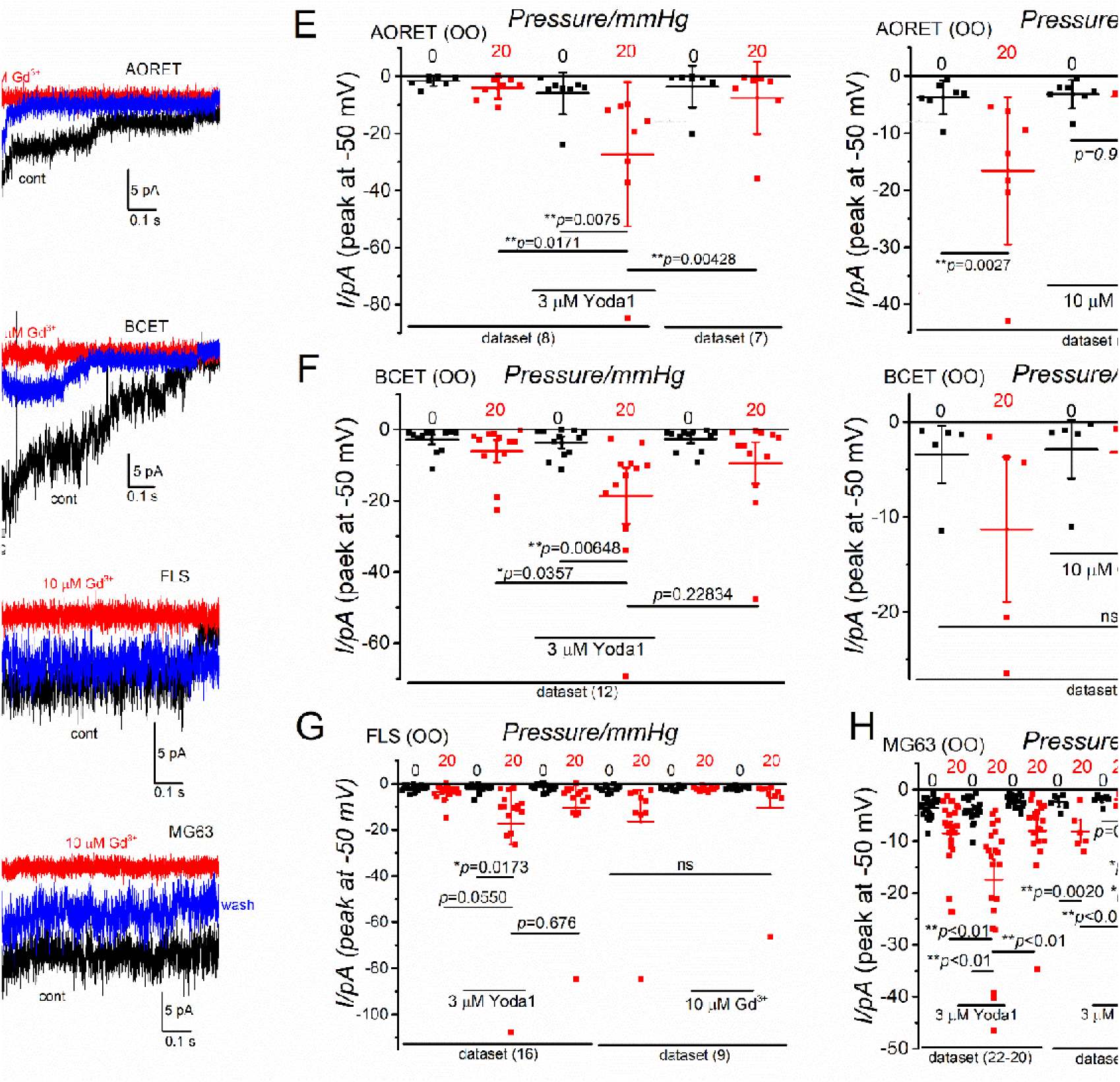
Effects of Yoda1 and Gd^3+^ on pressure-induced currents in human cells under the excised outside-out patch conditions. (A-D) Representative current traces recorded in the absence and presence of 20 mmHg positive pressure before (black), during (red), and after washout (blue) of 10 μM Gd^3+^ in AORET (A), BCET (B), FLS (C), and MG63 (D) cells. (E-H) Summary of peak current amplitudes measured in the absence of pressure (0 mmHg) and during application of 20 mmHg positive pressure in the absence or presence of 3 μM Yoda1 or 3-10 μM Gd. Statistical significance was assessed using Tukey’s post hoc test. The number of independent experiments is shown for each dataset. The ns indicate statistically no significance.

## 3. Discussion

In the present study, the author characterized the electrophysiological properties of endogenous human PIEZO1 (hPIEZO1) and compared them across four human cell types: human aortic endothelial cells (AORET), human brain capillary endothelial cells (BCET), human fibroblast-like synoviocytes (FLS), and human osteoblast-like MG63 cells (MG63). Endogenous hPIEZO1 currents in all four cell types were robustly activated by Yoda1 under both whole-cell and excised outside-out patch-clamp conditions. Furthermore, positive mechanical pressure applied to excised outside-out membrane patches effectively activated hPIEZO1-like currents, which were potentiated by Yoda1 and inhibited by Gd³. The unitary current amplitudes elicited by mechanical stimulation were also comparable to those elicited by Yoda1. These electrophysiological properties are broadly consistent with those previously observed for heterologously expressed hPIEZO1 in HEK cells (46.0 mmHg for p50, [25]). Collectively, the present findings demonstrate that endogenous hPIEZO1 exhibits broadly conserved electrophysiological properties across these four human cell types and support its role as a mechanosensor in diverse human tissues.

PIEZO1 is effectively activated by the synthetic agonist Yoda1 [9], which robustly stimulates hPIEZO1 and mPIEZO1 and has been reported to modulate PIEZO1 homologs in several other species, including rat, canine, crayfish, and *C-elegans* PIEZO1 [26–29]. In the present study, Yoda1 reversibly activated endogenous hPIEZO1 currents in all four human cell types under both whole-cell and excised outside-out patch-clamp conditions, with no clear evidence of cell type-specific differences under the experimental conditions employed. Three residues proposed to contribute to Yoda1 sensitivity and located between transmembrane helical units (THUs) 8 and 9 of mPIEZO1 (A1719, A2091, and A2094, [30]) are conserved in hPIEZO1 (A1718, A2075, and A2078). Mutation of A2091 to tryptophan in mPIEZO1 modestly reduces responses to high concentrations of Yoda1 [30], supporting a role for this region in Yoda1-dependent modulation of PIEZO1. Under our experimental conditions, endogenous hPIEZO1 currents were significantly potentiated by 3 μM Yoda1. This concentration is lower than the reported apparent EC_50_ of 26.6 μM for hPIEZO1 heterologously expressed in HEK cells [9]. In our previous study, however, HEK-hPIEZO1 exhibited an EC_50_ of 2.96 μM for Yoda1 [25]. Differences in experimental conditions may, at least in part, account for these apparent discrepancy in Yoda1 sensitivity. For example, solution flow during electrophysiological recordings may impose shear stress on the plasma membrane and thereby alter the mechanical state of PIEZO1, potentially facilitating Yoda1-induced activation. Consistent with this possibility, PIEZO1 in mouse vascular endothelial cells can be activated by Yoda1 at concentrations below 1 μM [31]. In addition, the lipophilic nature of Yoda1 [9] may influence its effective concentration within the membrane environment. Although, the precise structural determinants of Yoda1 binding and activation remains incompletely defined [32], the present findings further support Yoda1 as a useful pharmacological tool for identifying and modulating endogenous PIEZO1 activity in diverse human cell types.

The unitary current amplitudes of Yoda1-activatred channels were also comparable among the four cell types (Figure 4). Negatively charge residues located immediately above the extracellular fenestration, which contribute to efficient ion conduction in mPIEZO1 (2393-DEEED-2397), differ partially from the corresponding sequence in hPIEZO1 (2377-NEEAD-2381) [33], potentially contributing to species-dependent difference in ion permeation. In contrast, negative charge residues within the inner helix [33] that influence unitary conductance, Ca^2+^ permeability, and ruthenium red sensitivity in mPIEZO1 (E2487, E2495, E2496) are conserved in hPIEZO1 (E2461, E2469, and E2470). The comparable unitary current amplitudes observed among the four human cell types are therefore consistent with the presence of a common hPIEZO1 pore architecture, although the potential influence of the cellular membrane environment on channel permeation cannot be excluded.

PIEZO1 channels can be activated by diverse mechanical forces, including membrane stretch and shear stress generated by physiological fluid flow. Other mechanical inputs, including membrane indentation, changes in cell volume, and substrate stiffness can also modulate PIEZO1 activity. In the present study, positive mechanical pressure applied to excised outside-out membrane patches activated hPIEZO1-like currents in all four human cell types (Figure 2). Although pressure-induced currents tended to be larger in AORET and BMAET cells, the magnitude of the responses varied considerably among individual membrane patches in all four cell types (Figure 2 and Supplementary Fig.S1B). Differences in channel abundance within individual patches, patch geometry, membrane tension, or other local membrane properties may contribute to this variability. Indeed, relatively subtle differences in the mode and magnitude of mechanical stimulation can substantially alter PIEZO1 and PIEZO2 responses [34]. Conversely, channel density itself does not appear to determine the intrinsic mechanical sensitivity of PIEZO1 [35]. For datasets that could be satisfactorily fitted with the Boltzmann equation, the estimated midpoint pressure (p50) and slope coefficient (dp) did not differ significantly among the four cell types (Supplementary Table S1). However, not all pressure–response datasets could be adequately fitted, particularly in MG63 cells. Thus, although the present data do not provide evidence for major differences in pressure sensitivity among the four cell types, subtle cell type-dependent differences cannot be excluded. Importantly, mechanically activated currents were markedly potentiated by Yoda1 and substantially inhibited by Gd³ (Figures 3 and 5). Moreover, the unitary current amplitudes of mechanically activated single-channel events were comparable to those of Yoda1-activated events in each cell type (Figure 4 and Supplementary Table S2). Notably, PIEZO1 currents appeared to exhibit relatively slow inactivation kinetics across all four human cell types examined in this study. These observations may be attributable, at least in part, to differences in membrane lipid composition [36, 37] and/or curvature-dependent sorting [38]. Given that the auxiliary proteins MDFIC and MDFI (members of the MyoD family inhibitor protein family) have been shown to slow PIEZO channel inactivation [39], endogenous expression of these proteins may account, at least in part, for the relatively slow inactivation kinetics observed in these cells. Further studies combining a quantitative analysis of PIEZO1 inactivation kinetics with an assessment of endogenous MDFIC and MDFI expression levels would provide valuable insights into the molecular mechanisms underlying the observed inactivation properties. Taken together, these findings strongly support the conclusion that endogenous hPIEZO1 constitutes a major component of the mechanically activated currents detected in present four human cell types.

## 4. Materials and Methods

### 4.1. Reagents

The following reagents were used: 2-[5-[[(2,6-Dichlorophenyl)methyl]thio]-1,3,4-thiadiazol-2-yl]-pyrazine (Yoda1, Tocris Bioscience, Bristol, UK), and gadolinium chloride (Gd^3+^, Fujifilm, Osaka, Japan). Each reagent was dissolved in solvent recommended by the manufacturer and further diluted to the desired final concentration immediately before use.

### 4.2. Cell culture

Human osteoblast-like MG63 cells (MG63, American Type Culture Collection, Manassas, VA) was cultured in Dulbecco’s modified Eagle’s minimum (DMEM; Sigma-Aldrich, Tokyo, Japan) supplemented with 10% heat-inactivated fetal calf serum (FCS; Sigma-Aldrich), penicillin G (100 U/mL, Meiji Seika Pharma Co., Ltd., Tokyo, Japan), and streptomycin (100 μg/mL, Meiji Seika Pharma Co., Ltd.). Human aortic endothelial cells (AORET, DS PHARMA BIOMEDICAL, Osaka, Japan), human brain capillary endothelial cells (BCET, DS PHARMA BIOMEDICAL), and human fibroblast-like synoviocytes (FLS, Cell Applications, Inc., San Diego, USA) were maintained in cell-specific culture media with 10% heat-inactivated FCS, penicillin G (100 U/mL), and streptomycin (100 μg/mL). Cells were maintained under standard culture conditions until use.

### 4.3. Electrophysiological experiments

Whole-cell and excised outside-out patch-clamp experiments were performed as described previously [25, 40]. Patch pipettes with 3–5 MΩ resistance were used when filled with Cs^+^-based pipette solution, in which 110 mM Cs-aspartate, 30 mM CsCl, 1 mM MgCl_2_, 10 mM HEPES, 1 mM EGTA, and 2 mM Na_2_ATP, were contained and the pH was adjusted to 7.2 with CsOH. Membrane currents and voltage signals were recorded with an EPC-800 amplifier (HEKA, Lambrechit, Germany) and digitized at 10 kHz (PCI6229, National Instruments Japan, Tokyo, Japan). Whole-cell currents and outside-out single channel currents were acquired and analyzed using WinWCPV4.5 and WINEDR3.38, respectively, both developed by Dr. John Dempster (University of Strathclyde, UK). The liquid junction potential between the pipette and bath solutions (−10 mV) was corrected when the aspartate-rich pipette solution was used. For whole-cell recordings, voltage-ramp voltage protocols from -110 mV to +90 mV for 300 ms was applied every 5 s from a holding potential of −10 mV. For outside-out patch recordings, voltage-ramp protocols were applied every 5 or 10 s from a holding potential of −50 mV or −60 mV, as indicated. Leak currents was not subtracted from the recorded currents. The standard HEPES-buffered bathing solution (SBS) contained 137 mM NaCl, 5.9 mM KCl, 2.2 mM CaCl_2_, 1.2 mM MgCl_2_, 14 mM glucose, 10 mM HEPES, and the pH was adjusted to 7.4 with NaOH. All electrophysiological experiments were performed at 25 ± 1 °C with superfusion at 5 mL/min. Positive pressure was applied to excised outside-out membrane patches using a custom-built pressure-pulse system [25] consisting of a pressure pump (Linicon LV-435A; NITTO KOHKI Co., Ltd., Tokyo, Japan), an electrically controlled switching valve (VY1D00-6M5; SMC Corporation, Tokyo, Japan), and a pressure-monitoring amplifier (PA-011; Nihon Kohden, Tokyo, Japan). For construction of pressure-response curves (Figure 2), pressure-evoked current amplitudes were fitted in Origin using the Boltzmann equation.

I=Imin+(Imax-Imin)/(1+exp [(p-p50)/dp]),

where Imin and Imax represent the minimum and maximum current amplitudes, respectively, p represents the applied pressure, p50 represents the midpoint pressure, and dp represents the slope coefficient.

### 4.4. Statistical analyses

Data are presented as individual data points together with the mean ± standard deviation (SD), as indicated. Comparisons between two groups were performed using paired or unpaired two-tailed Student’s *t*-tests, as appropriate. Comparisons among multiple groups were performed using one-way analysis of variance (ANOVA) followed by Tukey’s post hoc multiple-comparisons test. Statistical analyses were performed using Origin J9.1 (LightStone Corp., Tokyo, Japan). A *p* value < 0.05 was considered statistically significant. The number of independent experiments (*n* or dataset) is indicated in the corresponding figures, tables, or figure legends.

### 4.5 Declaration of AI-assisted technology in the writing process

During the preparation of the work, the author used ChatGPT to improve the language and readability of the text. Following the use of this tool, the author carefully reviewed and edited the content as necessary and take full responsibility for the final content of the publication.

## 5. Conclusions

This study provides a direct electrophysiological comparison of endogenous hPIEZO1 currents across four distinct human cell types under comparable experimental conditions. Despite their different tissue origins, endogenous hPIEZO1 channels exhibited broadly similar responses to Yoda1 and mechanical stimulation, as well as comparable single-channel current amplitudes. These findings suggest that the fundamental electrophysiological properties of hPIEZO1 are largely conserved across these human cell types, whereas differences in channel expression, membrane composition, or other cellular factors may modulate the magnitude and kinetics of PIEZO1-mediated responses. Given the widespread expression of PIEZO1 throughout the human body, further studies aimed at defining tissue-specific regulation of PIEZO1 will be important for understanding its diverse physiological and pathophysiological roles and for evaluating the potential of PIEZO1 as a tissue-targeted therapeutic strategy.

## Supporting information

Supplementary Information revision1

## Supplementary Information

Supplementary information can be found.

## Acknowledgments

The author thanks Dr. J. Dempster (University of Strathclyde, UK) for developing the electrophysiology software (WinWCP4.5 and WinEDR3.38) and Drs H. Suzuki, A. Morita, and N. Hatano (Aichi-Gakuin University) for their technical support.

## Funding and additional information

This work was supported by Grants-in-Aid for Scientific Research to KM from Japanese Society of Promotion and Science (JSPS, 20K07599).

## Data availability

The data presented in this study are available on request from the corresponding author.

## Author Contributions

KM contributed intellectually, generated figures, analyzed and conceptualized the data, and performed the patch-clamp experiments supported. KM designed and initiated the project, acquired research funds, led and coordinated the project, interpreted the data, and wrote the paper.

