## Supplementary Information revision1 for "Electrophysiological Comparison of Mechanosensitive PIEZO1 Channel Currents across Different Human Cell Types"

*Revision_1*

Katsuhiko Muraki*

Laboratory of Cellular Pharmacology, School of Pharmacy, Aichi-Gakuin University, 1-100 Kusumoto, Chikusa, Nagoya 464-8650, Japan

Supplementary Table S1

| Cell type | n success fitting | mean of p50 (mmHg) | SD | mean of dp (slope) | SD | n total |
|---|---|---|---|---|---|---|
| AORET | 12 | 47.76704 | 19.5976 | 10.22299 | 5.38466 | 19 |
| BCET | 4 | 46.16249 | 9.37325 | 9.82229 | 4.73774 | 10 |
| FLS | 3 | 56.47224 | 6.71121 | 7.67775 | 2.54223 | 8 |
| MG63 | 4 | 34.78988 | 7.28848 | 9.09606 | 5.49127 | 23 |
| | | $p > 0.05$ | | $p > 0.05$ | | |

*Supplementary Table 1 Boltzmann function parameters in human cells.* Data are presented as the mean and standard deviation (SD) of p50 (mmHg) and dp (slope) parameters obtained by Boltzmann fitting of the pressure-induced currents in Figure 2. Each number of total and success fitting sample size is also denoted. Statistical significance was assessed using Tukey's post hoc test.

Supplementary Table S2

| Cell-type | Treatments | close | open1 | open2 | open3 | open4 | open5 |
|---|---|---|---|---|---|---|---|
| AORET | Yoda (3 μM) | Mean= -0.00948<br>SD= 0.05945 | Mean= -1.17226<br>SD= 0.21626 | Mean= -2.35962<br>SD= 0.314704 | Mean= -3.422<br>SD= 0.162635 | | |
| | 20mmHg | Mean= 0.023685<br>SD= 0.05945 | Mean= -1.25203<br>SD= 0.228065 | Mean= -2.46883<br>SD= 0.256211 | | | |
| BCET | Yoda (3 μM) | Mean= -0.05725<br>SD= 0.077976 | Mean= -1.2096<br>SD= 0.149888 | Mean= -2.28933<br>SD= 0.231368 | Mean= -3.531<br>SD= 0.161613 | | |
| | 20mmHg | Mean= 0.01003<br>SD= 0.045406 | Mean= -1.32078<br>SD= 0.208751 | Mean= -2.85675<br>SD= 0.240544 | | | |
| FLS | Yoda (3 μM) | Mean= -0.03321<br>SD= 0.031438 | Mean= -1.25812<br>SD= 0.204978 | Mean= -2.41988<br>SD= 0.219604 | Mean= -3.48975<br>SD= 0.280222 | | |
| | 20mmHg | Mean= 8.01E-04<br>SD= 0.042068 | Mean= -1.25812<br>SD= 0.158247 | Mean= -2.5045<br>SD= 0.226292 | | Mean= -4.5465<br>SD= 0.307801 | |
| MG63 | Yoda (3 μM) | Mean= -0.012<br>SD= 0.039681 | Mean= -1.30931<br>SD= 0.16304 | Mean= -2.554<br>SD= 0.218131 | Mean= -2.99667<br>SD= 0.136137 | Mean= -4.28867<br>SD= 0.246965 | Mean= -5.487<br>SD= 0.292527 |
| | 20mmHg | Mean= -0.01106<br>SD= 0.045623 | Mean= -1.2972<br>SD= 0.148639 | Mean= -2.643<br>SD= 0.215173 | Mean= -3.49133<br>SD= 0.214556 | | |

*Supplementary Table 2 Gaussian component parameters of unitary current amplitude in human cells.* Data are presented as the mean and standard deviation (SD) of parameters obtained by Gaussian fitting of the current amplitudes in Figure 4. Components with small sample sizes (n=2 and 3) were shown in underline.

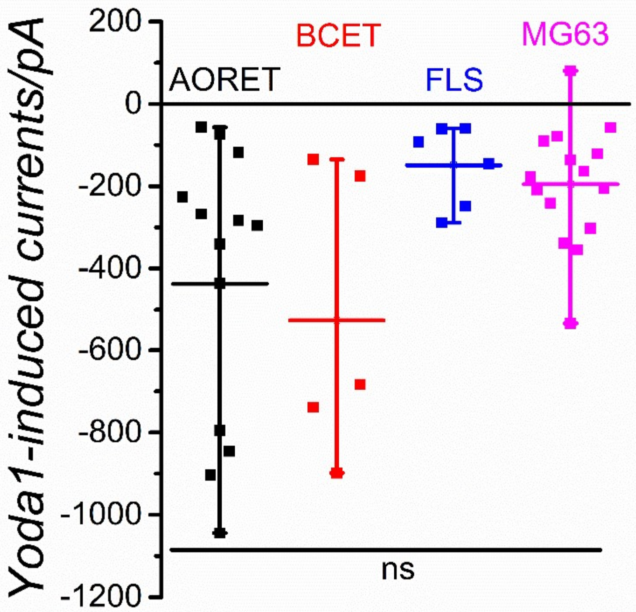


Supplementary Fig.S1A

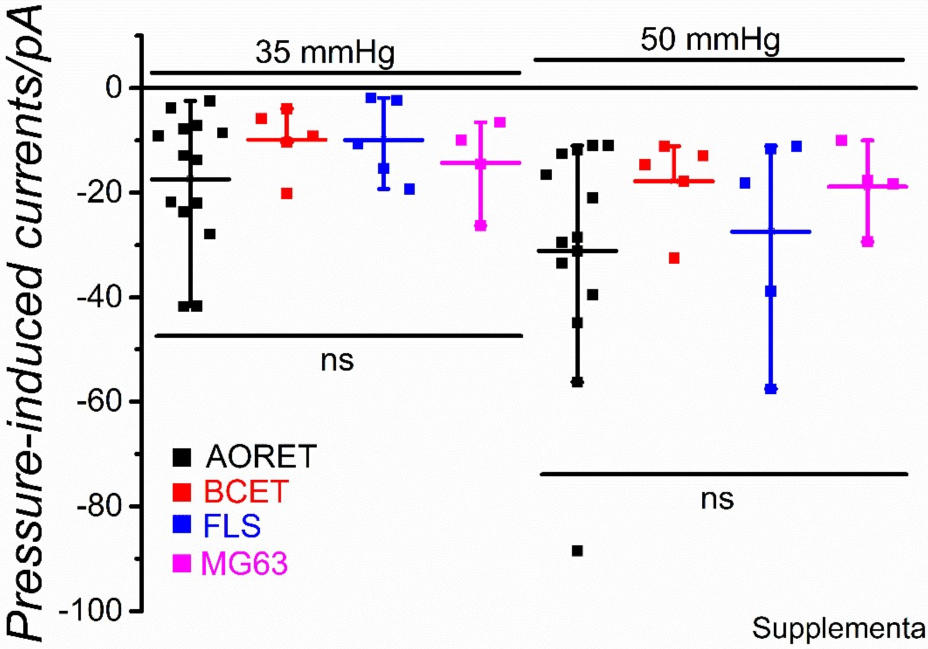


Supplementary Fig.S1B

*Supplementary Figure 1. Yoda1- and pressure-induced currents in human cells.* Peak amplitudes of currents induced by 3 μM Yoda1 (Fig.S1A) and 35 or 50 mmHg pressure (Fig.S1B) at -90 mV were summarized using the datasets shown in Figure 1 and 2, respectively. The numbers of independent experiments were 13, 5, 6, and 15 for AORET, BCET, FLS, and MG63 cells in Yoda1-induced currents, respectively, and 14, 5, 5, and 4 for AORET, BCET, FLS, and MG63 cells in 35 and 50 mmHg pressure-induced currents, respectively. Statistical significance was assessed using Tukey's post hoc test. The ns indicates statistically no significance.
